# Isolating structured salient variations in single-cell transcriptomic data with StrastiveVI

**DOI:** 10.1101/2023.10.06.561320

**Authors:** Wei Qiu, Ethan Weinberger, Su-In Lee

## Abstract

Single-cell RNA sequencing (scRNA-seq) has provided deeper insights into biological processes by highlighting differences at the cellular level. Within these single-cell omics measurements, researchers are often interested in identifying variations associated with a specific covariate. For instance, in aging research, it becomes vital to differentiate variations related to aging. To address this, we introduce StrastiveVI (<u>Str</u>uctured Contr<u>astive V</u>ariational Inference; https://github.com/suinleelab/StrastiveVI), which effectively separates the variations of interest from other dominant biological signals in scRNA-seq datasets. When deployed on aging and Alzheimer’s disease (AD) datasets, StrastiveVI efficiently isolates aging and AD-associated patterns, distinguishing them from dominant variations linked to sex, tissue, and cell type that are unrelated to aging or AD. In doing so, it underscores both well-known genes and potential novel genes related to aging or AD.

## 1 Introduction

When investigating various biological systems through single-cell omics measurements, it is often of interest to discern variations, such as changes in gene expression patterns, that are associated with a specific covariate of interest, as opposed to those that remain consistent regardless of the covariate. To illustrate, if we consider ‘chronological age’ as our covariate, we may want to distinguish variations linked to the aging processes from those that are unrelated to age.

Many recent works have proposed probabilistic latent variable models for the analysis of single-cell data [1–4]. However, a significant limitation of most of these models is their inability to effectively handle such analyses. This limitation arises because these models capture all the data’s variations using a *single* set of latent variables. Consequently, the learned representations by these models can mix together the variations of interest (e.g., related to age) with those that are not of interest.

Recent advancements in contrastive analysis [5–7] have addressed this issue, primarily in situations where each sample falls into one of two categories: a case group (e.g., cells subjected to drug exposure or CRISPR-induced genomic perturbations) or a corresponding control group (i.e., unperturbed cells). In such scenarios, the focus is on uncovering patterns that are overrepresented in the case cells compared to the control group. However, this framework has inherent limitations, as it is tailored to specific use cases. For instance, when attempting to disentangle age-related variations from aging-invariant relationships within single-cell RNA-seq data, we lack an explicit control group that can be employed to isolate the variations associated with aging.. Consequently, there is a pressing need for the development of new computational tools capable of addressing a broader array of contrastive research questions in this context.

As a first step towards this goal, here we introduce StrastiveVI (Structured Contrastive Variational Inference). StrastiveVI leverages previous advances in conditionally invariant representation learning [8] to model the variations underlying scRNA-seq data using two sets of latent variables: the first, called the *background variables*, are invariant to the given covariate of interest, while the second, called the *target variables*, capture variations related to the covariate of interest (e.g. age). We highlight StrastiveVI’s capabilities by applying it to scRNA-seq datasets related to aging and Alzheimer’s disease (AD) and find that it efficiently isolates aging and AD-specific signals from dominant biological signals that are unrelated to aging or AD.

In aging research, the advent of single-cell datasets designed for aging studies has prompted aging analysis at the cellular level [9–11]. However, tissue and cell type-related patterns often overshadowed aging signals. This complexity necessitated the creation of individual linear models for different tissue-cell type combinations, which not only consumed significant time but also risked missing globally significant aging genes. StrastiveVI tackles this challenge by effectively isolating aging-specific signals from the dominant tissue and cell type variations, enabling the identification of global aging variations across all tissues and cell types using a unified model.

## 2 Methods

### 2.1 The StrastiveVI generative process

For a target data point *x*_*n*_ we assume that each expression value *x*_*ng*_ for sample *n* and gene *g* is generated through the following process:

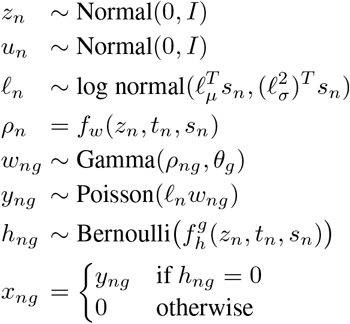

Here *z*_*n*_ and *u*_*n*_ denote two sets of latent variables that account for variations in scRNA-seq expression data. Specifically, *z*_*n*_ represents “background” variations that are invariant to the given target covariate of interest *t*_*n*_ (e.g. age), and which instead reflect variations due to a separate set of background covariates *b*_*n*_ that are not of interest for a given analysis (e.g. cell type). In contrast, *u*_*n*_ represents a set of “target” latent variables that reflect variations due to the target covariates and which are invariant with respect to the background covariates. We place a standard multivariate Gaussian prior on both sets of latent factors, as such a specification is computationally convenient for inference in the VAE framework [12]

To ensure that *z*_*n*_ and *u*_*n*_ indeed reflect their corresponding variations (i.e., background-covariate-related variations for *z*_*n*_ and target-covariate-related variations for *u*_*n*_), we employ the following two strategies. First, when optimizing the parameters of our generative model we include an additional loss term to ensure that a separate set of prediction neural networks can successfully recover the corresponding covariates from each set of latent variables: i.e., a background prediction network is trained to use *z*_*n*_ to predict values of the background covariates *b*_*n*_, while a target prediction network is trained to use *u*_*n*_ to predict the target covariates *t*_*n*_. Target covariates *t*_*n*_ and background covariates *b*_*n*_ can be binary, categorical, or continuous, and the exact loss function for the prediction networks thus depends on the nature of the covariate (e.g. cross-entropy loss is used for binary and categorical covariates, while mean squared error is used for continuous ones). Second, we employ the *d*-variable Hilbert-Schmidt Independence Criterion (*d*HSIC) [13–15, 8], a measure of the statistical dependence between random variables, to enforce independence between *z*_*n*_ and *u*_*n*_. The *d*HSIC between two variables is zero if and only if the variables are independent; thus, by minimizing the *d*HSIC we ensure that the target and background latent representations capture distinct biological signals.

Here *𝓁*_*μ*_ and 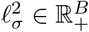, where *B* denotes the cardinality of an optional label denoting experimental batch, parameterize the prior for a latent RNA library size scaling factor on a log scale, and *s*_*n*_ is a *B*-dimensional one-hot vector encoding the batch label for each cell. For each batch, *𝓁*_*μ*_ and 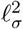 are set to the empirical mean and variance of the log library size. *ρ*_*ng*_ ℝ _+_ and shape *θ*_*g*_ ℝ _+_ parameterize our Gamma distribution with a mean-shape parameterization. Moreover, we note that *θ* _*g*_ can be viewed as a gene-specific inverse dispersion parameter for a negative binomial distribution, and we learn 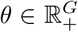 through variational inference. *f*_*w*_ and *f*_*g*_ are neural networks that transform the latent space and batch annotations to the original gene space, i.e. *f*_*w*_: ℝ ^*d*^ × {0, 1} ^*B*^ → ℝ ^*G*^ and similarly for *f*_*g*_, where *d* is the combined dimensionality of the concatenated target and background latent spaces. The outputs of the network *f*_*w*_ represent the mean proportion of transcripts expressed across all genes, and this constraint is enforced by using a softmax activation function in the last layer. That is, letting 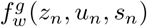 denote the entry in the output of *f*_*w*_ corresponding to gene *g*, we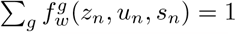 have. The output of the neural network *f*_*h*_ denotes whether a dropout event has occurred (i.e., a gene’s expression is read as zero due to technical factors rather than meaningful biological phenomena).

Our generative process closely follows that of contrastiveVI [7], but with the notable additions of prediction networks and the *d*HSIC penalty. While contrastiveVI’s modeling approach excels situations when explicit case and control groups of cells are available, it is not appropriate for scenarios without a clear control group. By incorporating target covariates *t*_*n*_ and background covariates *b*_*n*_, and introducing prediction networks for both, StrastiveVI can effectively isolate the target-covariate-induced variations even without an explicit group of corresponding control cells.

### 2.2 Inference with StrastiveVI

We cannot compute the StrastiveVI posterior distribution using Bayes’ rule as the integrals required to compute the model evidence *p*(*x*_*n*_ | *s*_*n*_) are analytically intractable. As such, we instead approximate our posterior distribution using variational inference [16]. We approximate our posterior with a distribution factorized as follows:

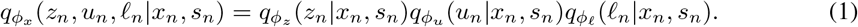

Here *ϕ*_*x*_ denotes a set of learned weights used to infer the parameters of our approximate posterior. Based on our factorization of the posterior in Eq. 1, we can divide our full set of parameters *ϕ*_*x*_ into disjoint subsets *ϕ*_𝓏_, *ϕ*_*u*_ and *ϕ*_𝓁_ for inferring the parameters of the distributions of *z, u* and 𝓁 respectively. As in the VAE framework [12], we approximate the posterior for each set of latent variables via a deep neural network that takes in expression levels as input and returns the parameters of its corresponding approximate posterior distribution. Moreover, we note that each factor in the posterior approximation shares the same family as its respective prior distribution (e.g. *q*(*z*_*n*_ | *x*_*n*_, s_*n*_) follows a normal distribution). By marginalizing out *w*_*ng*_, *h*_*ng*_, and *y*_*ng*_, we can simplify our likelihood yielding *p*_𝒱_ (*x*_*ng*_ | *z*_*n*_, u_*n*_, s_*n*_, 𝓁_*n*_), which has a closed form of a zero-inflated negative binomial (ZINB) distribution and where *v* denotes the parameters of our generative model. We implement our generative model with deep neural networks as done for our approximate posterior distributions. For Equation 1 we derive the following variational lower bound:

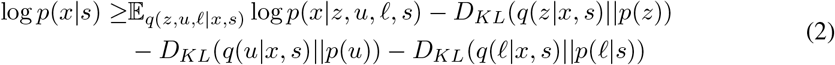

We optimize the parameters of our generative model, inference networks, and prediction networks in a joint manner using stochastic gradient descent. The objective is to minimize a composite loss comprising the negative ELBO from Eq. 2, prediction losses for *t*_*n*_ and *b*_*n*_, and the *d*HSIC. The neural networks underpinning both the variational and generative distributions are feedforward with standard activation functions.

## 3 Results

To assess the performance of StrastiveVI, we first utilize it to isolate variations related to aging from aging-invariant relations. Given that chronological age is a continuous variable and no “control” value for age exists, using previously proposed contrastive methods is not possible for this task. Moreover, to demonstrate StrastiveVI’s ability to extract discrete variations, we apply it to isolate variations associated with Alzheimer’s disease.

### 3.1 Isolating aging-related variations

To assess StrastiveVI’s performance in isolating variations related to aging, we use it on the Tabula Muris Senis (TMS) droplet dataset [17], a large expert-curated scRNA-seq dataset. This dataset includes cells from mice (mus musculus) across different ages, such as 1m, 3m, 18m, 21m, 24m, and 30m, making it an invaluable resource for studying aging at the single-cell level across various tissues and cell types. We focus on the 20 most common cell types. In total, the data we use contains 124,373 cells taken from 14 different tissues and organs. We apply StrastiveVI to extract variations related to aging. Our goal is to isolate aging signals across all sexes, tissues, and cell types. Supplementary Figure 1 illustrates that the raw data’s dominant biological signals include sex, tissue, and cell type. Therefore, we treat chronological age as the target covariate (*t*), with sex, tissue, and cell type serving as background covariates (*b*). We split the dataset into 64% training, 16% validation, and 20% test sets. An effective model should learn a target latent space that captures aging-related variations while retaining other biological patterns unrelated to aging in the background latent space.

**Figure 1.**
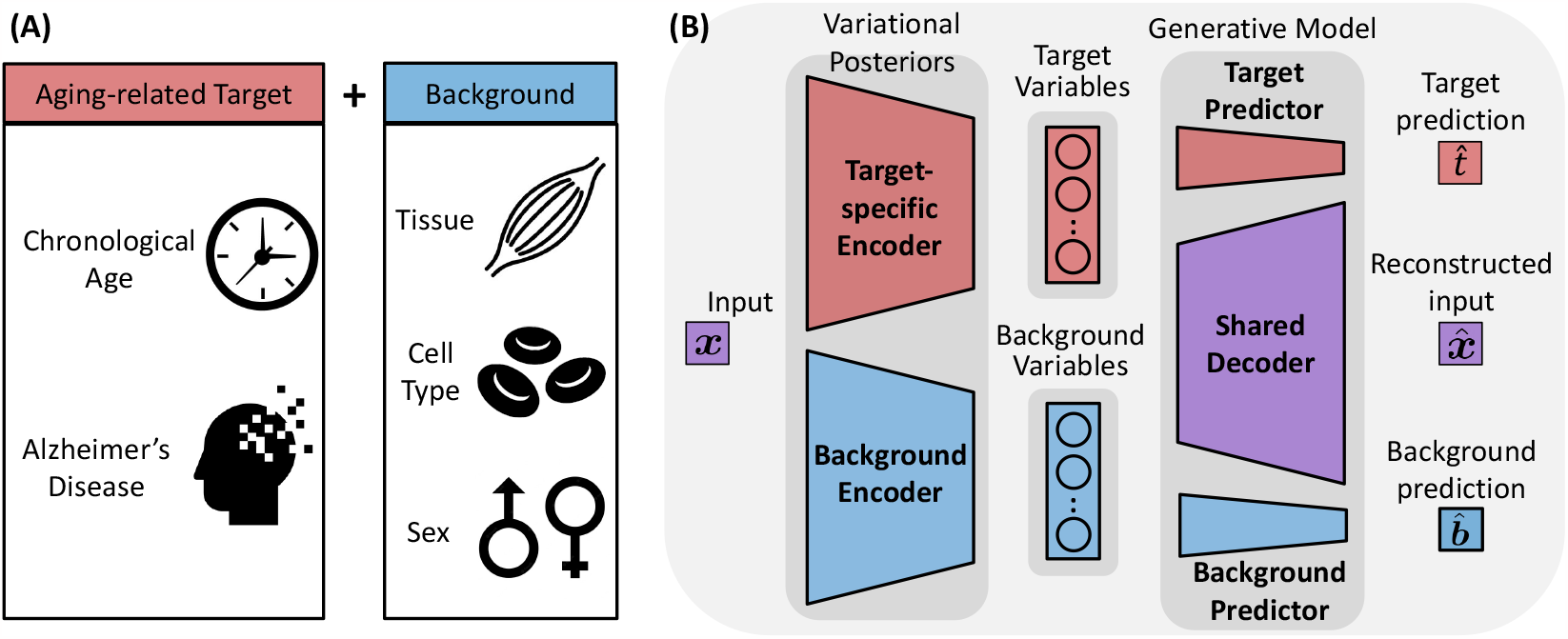
For a given dataset, alongside its target covariate *t* (covariate of interest) and background covariates *b*, StrastiveVI separates the target variations and the dominant background variations. ((A) We apply StrastiveVI to identify aging-related variations. We concentrate on general aging trends by taking chronological age as the target covariate and on aging-related diseases, specifically Alzheimer’s disease, as the target covariate. The dominant biological variations (background covariates), which are not of interest, comprise tissue, cell type, and sex. (B) An overview of the StrastiveVI structure: StrastiveVI consists of two encoder networks. The target-specific encoder embeds a cell into the target latent space, and the background encoder embeds a cell into the background latent space. The model incorporates a target predictor for predicting the target covariate from the target variables and a background predictor for predicting background covariates from the background variables. This setup ensures that the target-specific encoder isolates variations related to the target covariate, and the background encoder captures other biological variances. The *d*-variable Hilbert-Schmidt Independence Criterion (*d*HSIC) penalty is applied to encourage independence between the target and background variables. Both target and background latent cell representations are then transformed back to the original gene expression space using a decoder network.

Figure 2A shows that the target latent space successfully separates aging-related signals, as seen by the clear age trend. In Figures 2B-D, cells from different sexes, tissues, and cell types are mixed together, indicating that the aging signals identified by the target latent variables are consistent across these different groups. Figures 2E,G,H suggest that the other biological signals about cell type, sex, and tissue that are unrelated to aging are captured in the background variables. Meanwhile, the aging signals in the background variables, as seen in Figure 2F, are subtle. From this, we can conclude that our model effectively isolates age-related variations in the target latent variables while preserving other critical biological signals in the background. This separation enhances our ability to gain a clear understanding of how aging impacts cells, free from interference from other factors.

**Figure 2.**
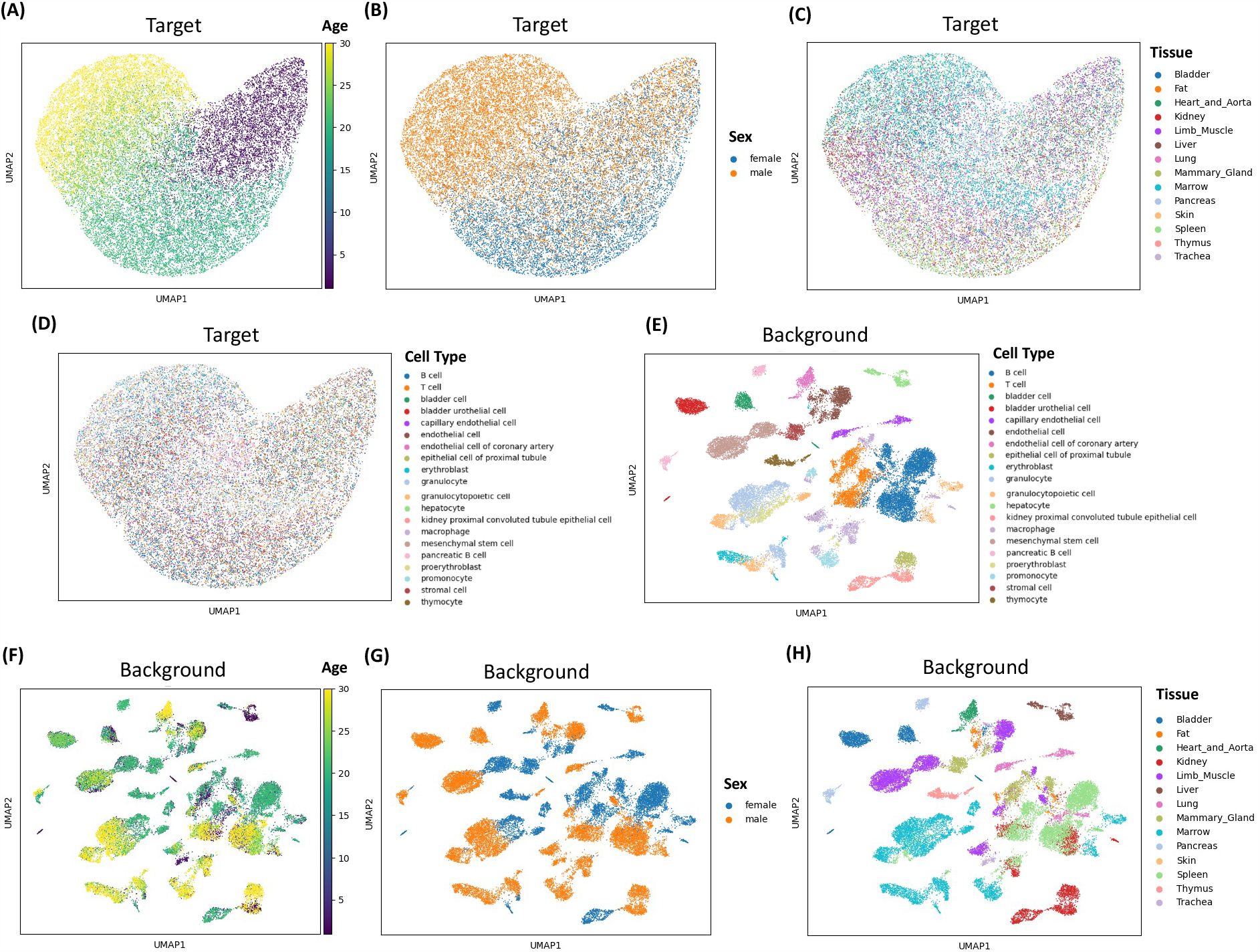
StrastiveVI isolates aging-related variations. (A)–(D), UMAP plots of StrastiveVI’s target latent representations, colored by chronological age (A), sex (B), tissue (C), and cell type (D). (E)–(H), UMAP plots of StrastiveVI’s background latent representations, colored by cell type (E), chronological age (F), sex (G), and tissue (H).

Moreover, we assess the target latent variables’ capability in age prediction. To do this, we employ a fully connected neural network that uses the target latent representations as input and is trained to predict chronological age. The model shows strong performance, with a Pearson’s *r* value of 0.91. Figure 3 underscores that cells from subjects of the same chronological age have predicted ages that span a range. This variation indicates the model’s capability to discern subtle differences in aging between cells. We further apply the Integrated Gradient, an attribution method for understanding feature importance in neural networks, on the target latent variable encoder and the age prediction model [18] to identify the important aging genes. Among the top 100 genes identified, 57 were already documented in GenAge [19] as associated with aging. This substantial overlap reinforces our model’s effectiveness in isolating aging-related signals. The alignment with GenAge also suggests that the remaining important genes we identify could be promising candidates for further exploration regarding their involvement in aging.

**Figure 3.**
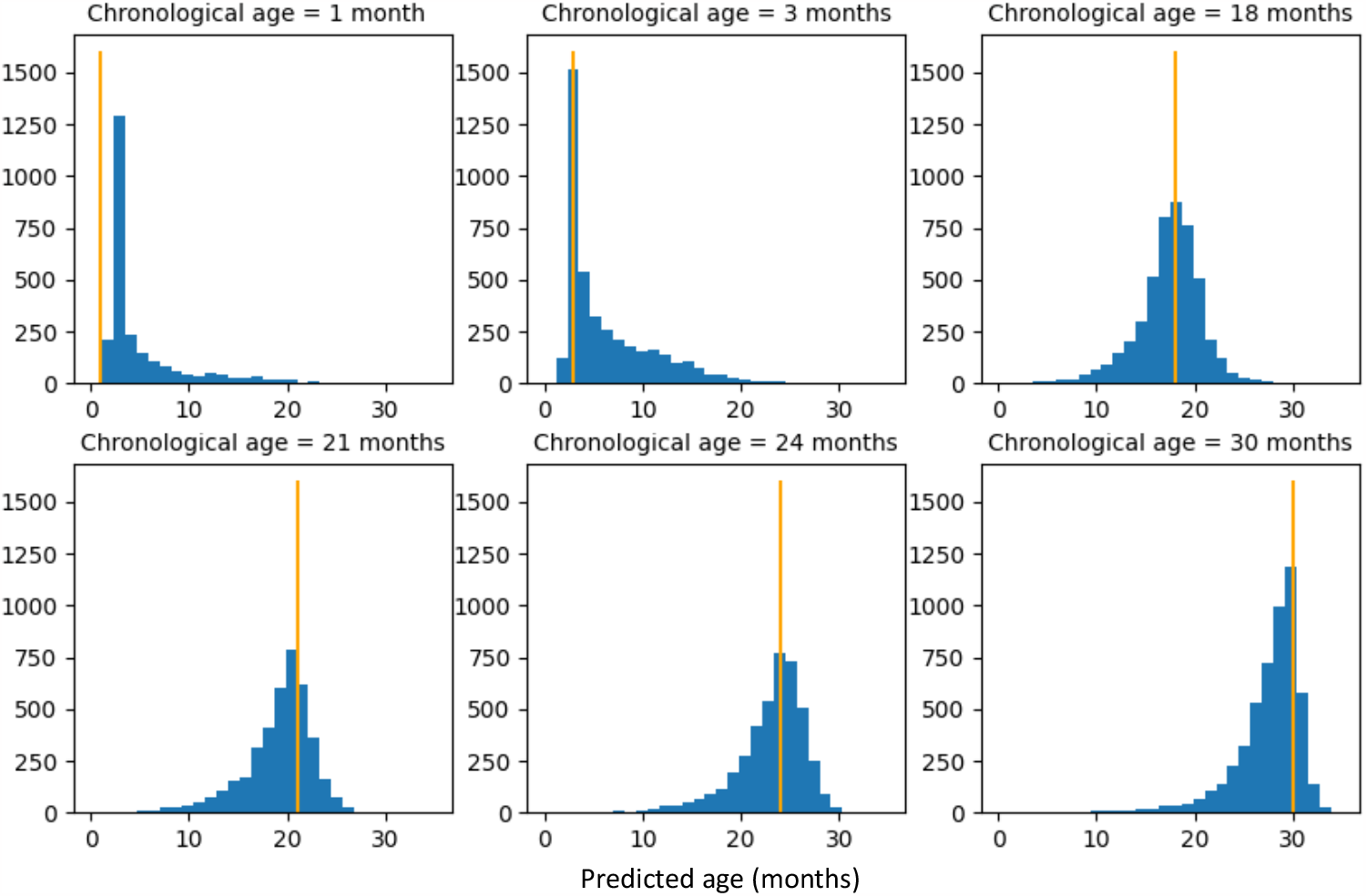
Histograms of predicted ages for cells from subjects with the same chronological age.

### 3.2 Isolating Alzheimer’s disease-related variations

We also evaluate our StrastiveVI model on Alzheimer’s disease (AD), an aging-related condition. We use a scRNA-seq dataset derived from entorhinal cortex samples from both control and AD brains, with six samples in each group [20]. This dataset comprises 13,214 cells from six cell types. Supplementary Figure 2 indicates that the main biological signals in the raw data relate to sex and cell type. As a result, we treated the disease condition (either AD or control) as the target covariate (*t*), while considering sex and cell type as background covariates (*b*). We divide the dataset into 64% for training, 16% for validation, and 20% for testing. Through training the StrastiveVI model, our goal is to isolate AD-related variations across all sexes and cell types.

Figure 4A demonstrates that the target latent space clearly separates cells from AD patients and those from control subjects. In Figures 4B-C, cells from different sexes and cell types mix together, suggesting that the AD-related variations identified by the target latent variables are consistent across these groups. Figures 4E-F show that signals related to cell type and sex are mainly captured by the background variables. Also, the AD-related signals in these background variables, as shown in Figure 4D, are barely discernible when compared to Supplementary Figure 2A. The separation between AD-related signals and other biological factors demonstrates the model’s capability to accurately identify patterns specific to Alzheimer’s disease.

**Figure 4.**
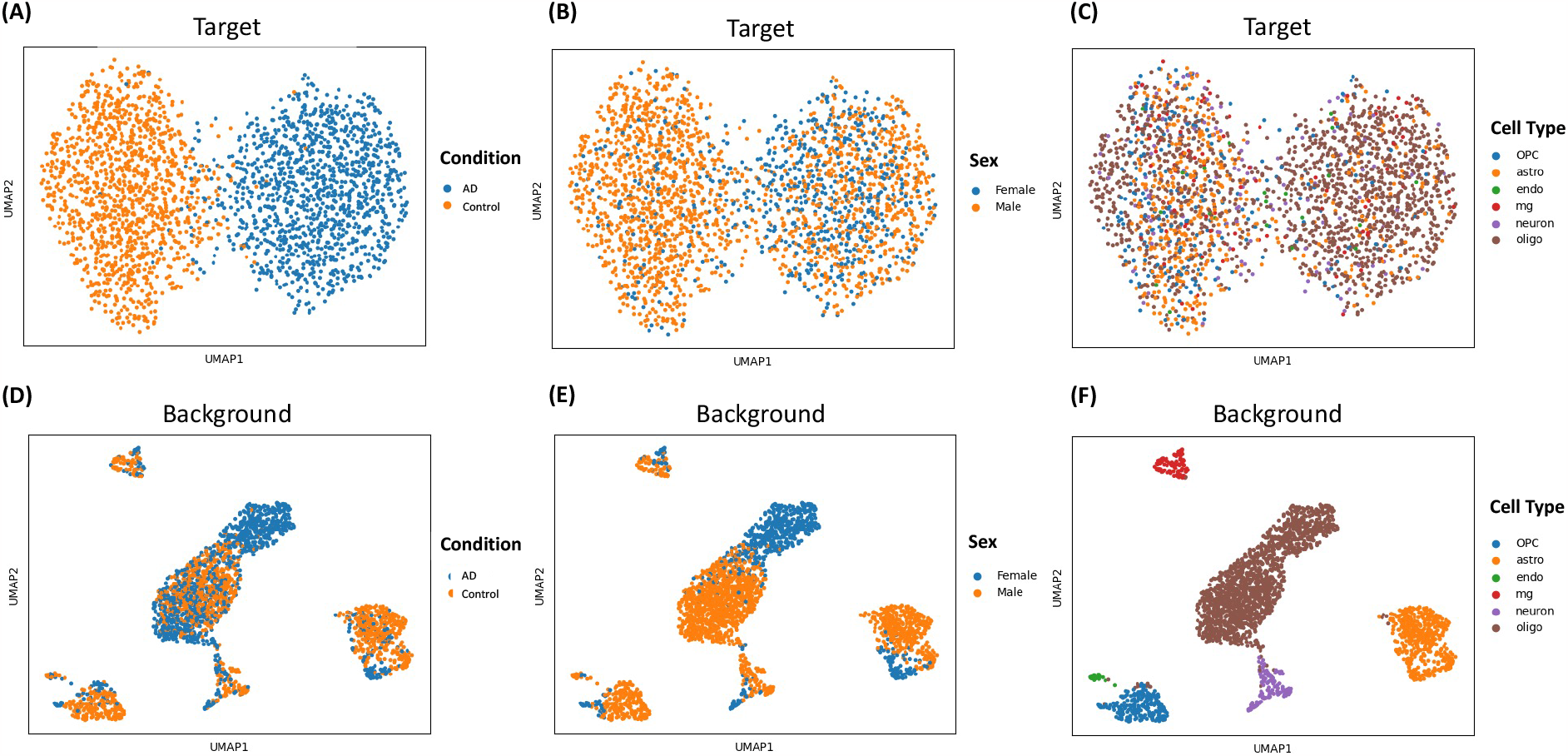
StrastiveVI isolates Alzheimer’s disease-related variations. (A)–(C), UMAP plots of StrastiveVI’s target latent representations, colored by disease condition (A), sex (B), and cell type (C). (D)–(F), UMAP plots of StrastiveVI’s background latent representations, colored by disease condition (D), sex (E), and cell type (F).

To better understand which genes drove this separation, we apply Hotspot [21], a tool that ranks genes based on spatial autocorrelation in relation to cell-cell similarity metrics, on the target latent space. Figure 5 displays 6 of the top 20 key genes. Their expression levels differ between AD and control cells, and some have been reported to be associated with AD in previous research [22–24]. The alignment of our model’s results with prior research not only highlights the significant roles these genes might play in AD progression but also underscores the effectiveness of StrastiveVI in extracting relevant biological insights.

**Figure 5.**
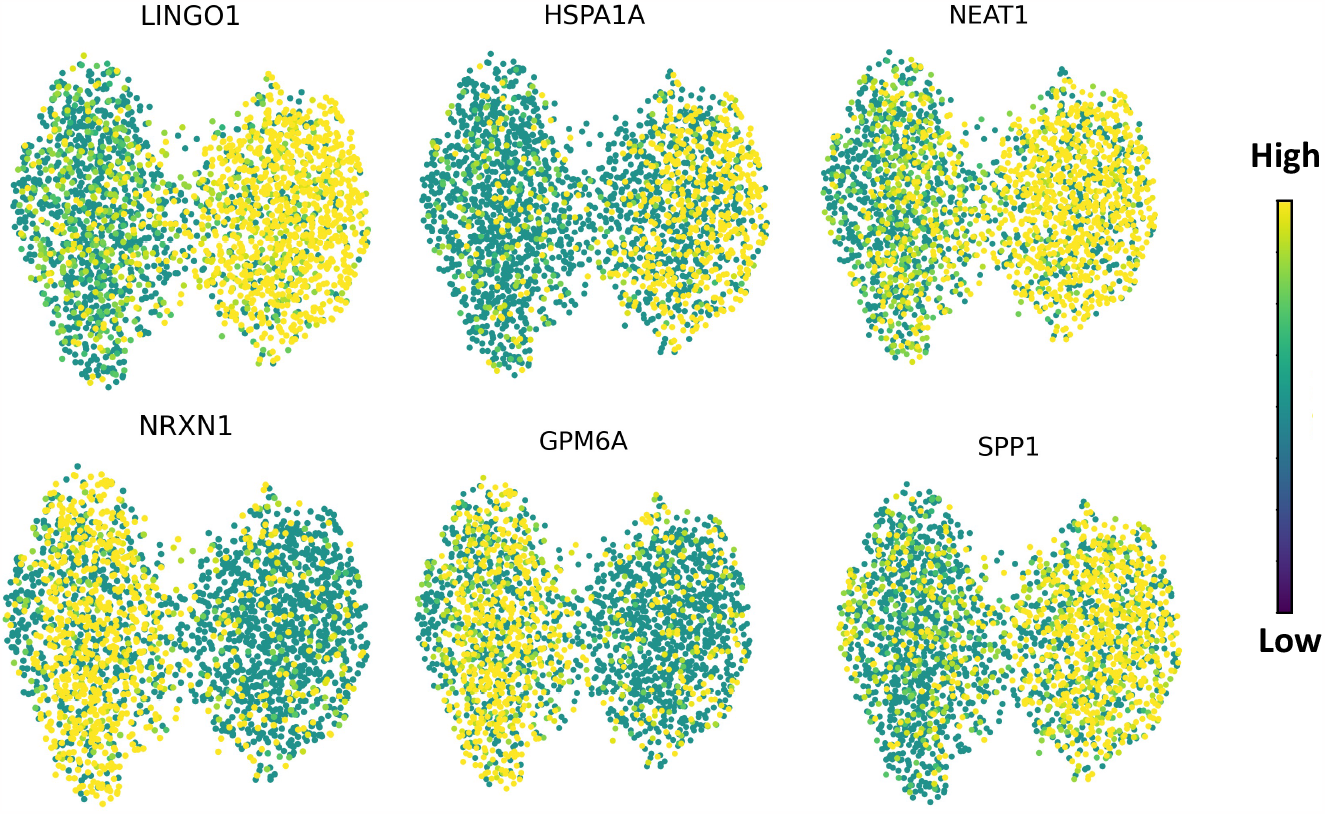
UMAP plots of StrastiveVI’s target latent representations, colored by expression levels of 6 genes returned by Hotspot.

## 4 Discussion

StrastiveVI effectively differentiates aging and Alzheimer’s disease patterns from other inherent biological signals (i.e., sex, cell type, and tissue). The distinct age trends and discernible AD conditions in the target latent space highlight the model’s proficiency in isolating these variations of interest. Simultaneously, other biology variations are allocated to the background space. Through this separation, StrastiveVI offers nuanced and profound insights into both the aging and AD processes, effectively accounting for and distinguishing other biological influencers. In our findings, the genes that emerged as vital in the target latent space align with those highlighted in earlier studies, emphasizing the effectiveness of StrastiveVI in recognizing key genes associated with aging and AD. Moreover, while our models confirm some known genetic markers, their strength lies in potentially revealing new insights and highlighting previously overlooked genes that may play crucial roles in the complex process of aging and AD.

StrastiveVI offers greater flexibility and broader applicability than contrastiveVI. contrastiveVI needs both a case group and a matching control group to pinpoint specific variations. However, in many scenarios, like the study of aging-related variations, identifying an explicit control group can be challenging. StrastiveVI’s design overcomes this by treating all cells as the case group and eliminating the necessity for a control group through the direct prediction of target covariates. Additionally, the introduction of background prediction networks in StrastiveVI helps in separating dominant biological patterns from the target space. Notably, StrastiveVI can process target and background covariates regardless of their type: binary, categorical, continuous, or graph-structured labels. This flexibility opens doors for deeper analyses and potential discoveries in various research areas.

Earlier studies leveraging transcriptomics data for aging analysis predominantly focused on bulk data [25–27]. With the advent of extensive scRNA-seq datasets specifically designed for aging research [17, 28, 29], researchers began exploring aging at the single-cell level [9–11]. A significant challenge is that biological patterns related to tissue and cell type often overshadow the subtler signals associated with aging. Consequently, many studies using single-cell data for aging analysis have constructed individual linear models for distinct tissue-cell type combinations which could be labor-intensive and can encounter issues with limited cell sizes for specific tissue-cell pairings. Most importantly, this approach might miss out on identifying genes associated with global aging processes that are consistent across different tissues and cell types. In this context, StrastiveVI emerges as a transformative solution. Its design enables us to effectively disentangle aging-related signals from the dominant tissue and cell type variations, capturing global aging patterns with a single model. By bypassing the need for separate models for each tissue-cell type combination, StrastiveVI offers a more streamlined and holistic approach to understanding the aging process at the single-cell level.

As we move forward, there are several avenues of exploration and validation. A critical next step is the introduction of quantitative metrics that allow for a more objective and detailed comparison of StrastiveVI with baseline models undertaking contrastive analysis. Additionally, embracing explainable methodologies for unsupervised models [30] could provide richer insights into the patterns within the target embedding space, possibly offering a more nuanced understanding than methods like Integrated Gradient, traditionally tailored for supervised models. While our computational tools and predictions provide significant insights, the empirical validation that wet labs offer remains unparalleled for biological validations. Rigorous lab experiments are essential to substantiate the relevance of genes that StrastiveVI associates with aging and AD. Such endeavors not only reaffirm our hypotheses but may also illuminate novel therapeutic opportunities in the domains of aging and AD.

StrastiveVI’s potential extends far beyond its current applications. One of its notable uses could be in determining a single-cell level biological age that’s consistent across different tissue and cell types. Such a unified metric would enhance our ability to assess cellular aging interventions with a finer resolution. Furthermore, the adaptable nature of StrastiveVI makes it well-suited for various challenges. In oncology, StrastiveVI can be utilized for pan-cancer analyses, using cancer types as target covariates, thereby discerning signals that are common across different cancers. In the realm of spatial transcriptomics, the model can be expertly applied to interpret patterns by considering spatial graphs as target covariates. These potential applications underscore the versatility of StrastiveVI and its potential role in shaping the future of cellular and molecular research.

