## Supplementary Material for "Isolating structured salient variations in single-cell transcriptomic data with StrastiveVI"

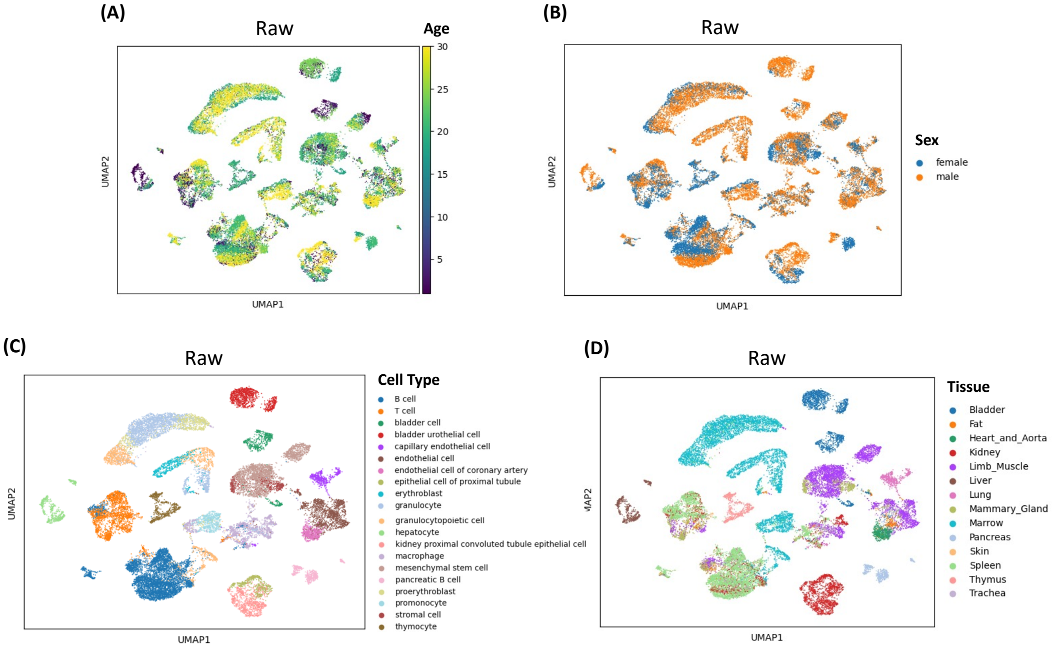


Supplementary Figure 1: (A)–(D), UMAP plots of the raw data for aging analysis colored by chronological age (A), sex (B), cell type (C), and tissue (D).

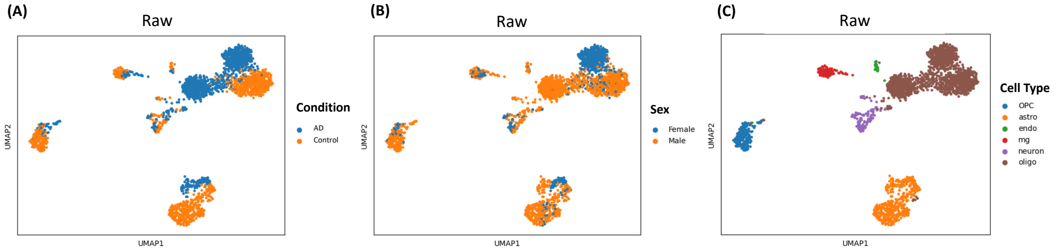


Supplementary Figure 2: (A)–(C), UMAP plots of the raw data for AD analysis colored by disease condition (A), sex (B), and cell type (C).
